# Age-dependent extinction and the neutral theory of biodiversity

**DOI:** 10.1101/2023.08.23.554417

**Authors:** James G. Saulsbury, C. Tomomi Parins-Fukuchi, Connor J. Wilson, Trond Reitan, Lee Hsiang Liow

## Abstract

Red Queen (RQ) theory states that adaptation does not protect species from extinction because their competitors are continually adapting alongside them. RQ was founded on the apparent independence of extinction risk and fossil taxon age, but analytical developments have since demonstrated that age-dependent extinction is widespread, usually most intense among young species. Here we develop ecological neutral theory as a general framework for modeling fossil species survivorship under incomplete sampling. We show that it provides excellent fit to a high-resolution dataset of species durations for Paleozoic zooplankton, and more broadly can account for age-dependent extinction seen throughout the fossil record. Unlike widely used alternative models, the neutral model has parameters with biological meaning, thereby generating testable hypotheses on changes in ancient ecosystems. The success of this approach suggests novel interpretations of mass extinctions and of scaling in eco-evolutionary systems. Intense extinction among young species does not necessarily refute RQ or require a special explanation, but can instead be parsimoniously explained by neutral dynamics operating across species regardless of age.

**Significance Statement:** Red Queen theory predicts that competition among species should cause extinction risk to be independent of species age, but recent analyses have refuted this central prediction. To fill the resulting theoretical vacuum, we used ecological neutral theory to build a model of the lifespans of incompletely sampled species evolving under zero-sum competition. This model predicts survivorship among fossil zooplankton with surprising accuracy and accounts for empirical deviations from the predictions of Red Queen more generally. A neutral model of background extinction allows for interpreting survivorship curves in terms of biological process, suggests a novel understanding of mass extinctions, and supports a role for competition in extinction.

## Introduction

How does extinction risk change over the lifetime of a species? This question motivated one of paleontology’s most significant contributions to evolutionary theory in Van Valen’s Red Queen (RQ) hypothesis, a model of evolutionary turnover in which species compete for limited resources against an ever-evolving set of competitors (1–4). This theory was first developed to explain the observation that extinction in fossil taxa is apparently independent of taxon age. A species never becomes buffered against extinction by adaptation because its competitors are constantly adapting alongside it (1). Species survivorship can thus speak to weighty macroevolutionary questions, but the evidence for age-independent extinction in the fossil record no longer seems firm. Improvements in statistical survivorship analysis and in paleontological data compilations indicate instead that age-independent extinction is the exception rather than the norm, with extinction typically most intense among young taxa (2, 5). For example, a recent survey found negative relationships between species age and extinction risk in 25 of 33 clades, with only 3 clades showing age-independence (6). Age is one of the most universal characteristics of lineages, so very general insights could come from understanding the “aging process” in fossil species. Nevertheless, the predominance of age-dependent extinction in the fossil record lacks a strong theoretical explanation.

A promising but underexplored approach to extinction is the neutral theory of biodiversity (NT) (7), a general model of the abundance dynamics of species in a community (Fig. 1). This model is neutral because individuals are competitively identical regardless of species identity; the success or failure of species is therefore stochastic, determined solely by zero-sum ecological drift. NT thus dispenses with some features of biological interest like ecologically determined variation in fitness (8), but the resulting simplicity allows the theory to generate testable and often successful predictions across a broad array of ecological and evolutionary variables, including species abundances, species-area relationships, and phylogenetic tree shape (9). Moreover, NT can be useful even when it fails, working as a null hypothesis for identifying non-neutral phenomena (10). Finally, NT has deep similarities with RQ (11), and both could potentially explain species survivorship as the result of general laws applying to all species at all ages: namely, zero-sum interactions among competitively equivalent ecological entities (11). However, the predictions of NT for age-dependent extinction in the fossil record are unexplored.

**Figure 1.**
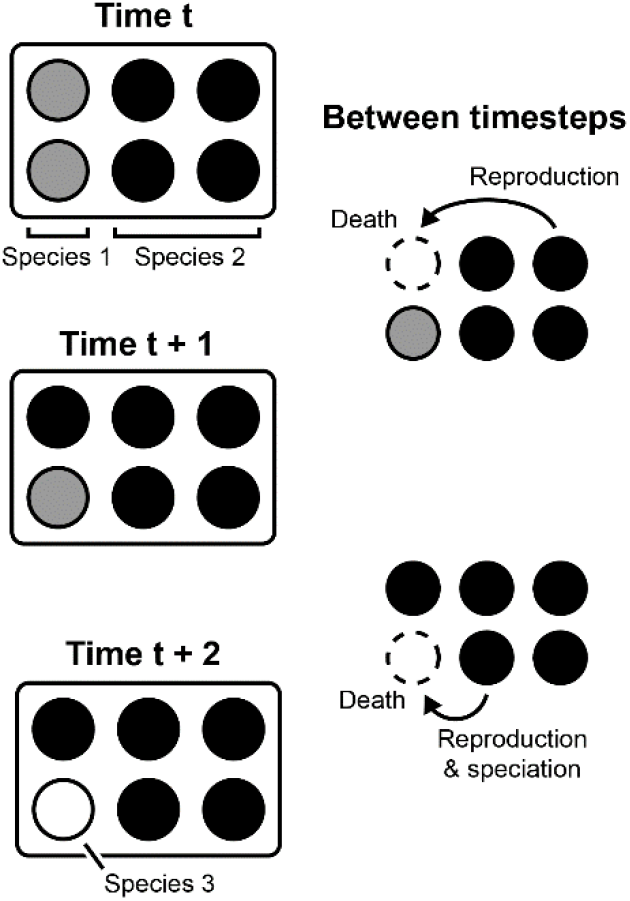
Extinction and speciation in a neutral community. Illustrated community includes six individuals belonging initially to two species. In each timestep, a random individual dies, a random surviving individual reproduces itself, and the new individual has a chance of becoming a new species.

We developed NT to investigate the relationship between species age and extinction rate under incomplete sampling, and used it to study an exceptional dataset of precise [20–100ky resolution (12)] stratigraphic ranges for 1794 species of Ordovician-Silurian graptoloids (13–15), an extinct clade of planktonic colonial suspension feeders. This dataset includes a mass extinction event and contrasting Ordovician and Silurian extinction regimes linked to changes in global climate (13). We derived expressions for the probability of observing any given species duration in NT under incomplete sampling. We then evaluated relative and absolute fit of these models to survivorship data using maximum likelihood estimates of community size *J*, speciation rate *v*, and sampling rate *s*, considering the full dataset as well as subsets of species originating during the Ordovician [481–447 Ma; excludes the late Katian and Hirnantian following (13)], Silurian (447– 419 Ma, including late Katian and Hirnantian), and immediately before the Late Ordovician Mass Extinction (450–448 Ma). NT was compared with two survivorship models that are not grounded in biological principles but are commonly used because of their simplicity: the exponential distribution, representing age-independent extinction, and the Weibull distribution, a generalization of the exponential in which extinction risk can increase or decay toward zero with species age. Our findings represent a novel test of ecological theory with fossil data, yield insights into evolutionary turnover in the marine plankton, and help put species survivorship on a solid theoretical foundation.

## Results

In NT, extinction dynamics emerge naturally from abundance dynamics (Fig. 2; Eqn. 6). Species that do not go extinct early on tend to increase from low starting abundance, causing extinction risk initially to decline with age (Fig. 2A-B). However, abundance of survivors eventually converges on a conditional stationary distribution (20), causing extinction rate to become age-independent among old species. The result is a two-phased survivorship curve. Increasing abundance with species age also gives older species a greater chance of being sampled (Fig. S1). The three free parameters of NT under incomplete sampling have straightforward effects on survivorship curve shape (Fig. 2C). Speciation rate controls the extinction rate in old species and community size determines the age at which species reach constant extinction risk. Lowering the sampling rate increases the bias against sampling very young species, such that survivorship appears more age-independent. Incorporating sampling is necessary to achieve good fit to empirical data because large neutral communities have an excess of rare, ephemeral species—a problem which has been solved in a different context by introducing alternate modes of speciation into NT (16). Despite its flexibility (Fig. 2C), NT is falsifiable, as it can only produce initially concave-up survivorship curves that eventually flatten out.

**Figure 2.**
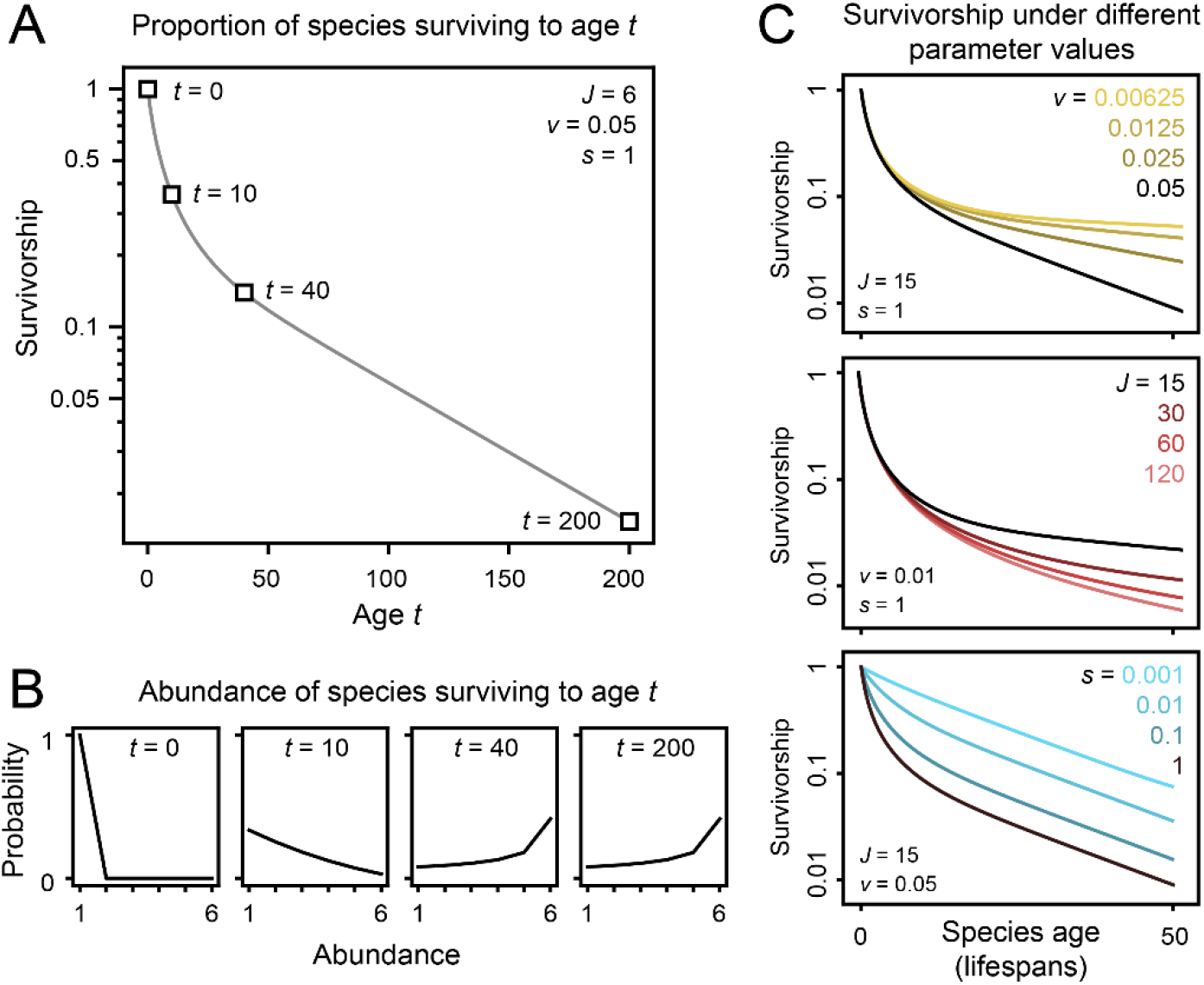
Survivorship in NT. (A) A survivorship curve, or semilog plot of species age *t* versus the probability that a species survives to age t. (B) Probability of a species having some abundance, given survival to age t. (C) The range of survivorship curves expected in NT under different values of speciation rate *v*, community size *J*, and per-individual per-timestep sampling rate *s*. Species age is given in lifespans, the expected number of timesteps until an individual dies.

NT successfully models empirical species survivorship, both relative to alternative models and in absolute terms. Empirical survivorship curves among graptoloids are known to have a characteristic shape, with intense extinction among very young (<0.2 My duration) species and low, age-independent risk in older species (16). A Weibull model with decreasing extinction risk is a better model than the exponential, but neither matches the two-phased structure of the empirical curve (Fig. 3A). Conversely, NT closely matches the data (Fig. 3B), reflected by decisive support from BIC (ΔBIC=1068 for exponential and 165 for Weibull; Table S1). BIC comparisons overwhelmingly support NT for Ordovician and Silurian subsets of the data as well (Tables S2-S3, Figs. S3-S4). NT fits imply the existence of many unsampled rare species: in the incompletely sampled model, ∼65% of species survive until the age-independent phase of the survivorship curve (Fig. 3B), whereas only ∼6% of species do so in a version of the same model with perfect sampling. Finally, Kolmogorov-Smirnov tests reveal NT is close to the empirical distribution, deviating from the empirical survivorship curve by a maximum of just 3% (0.05>p>0.02) for the full dataset, compared with 20% (p<0.001) for the exponential and 9% (p<0.001) for the Weibull. We obtain similar results for Ordovician and Silurian subsets (Table S5).

**Figure 3.**
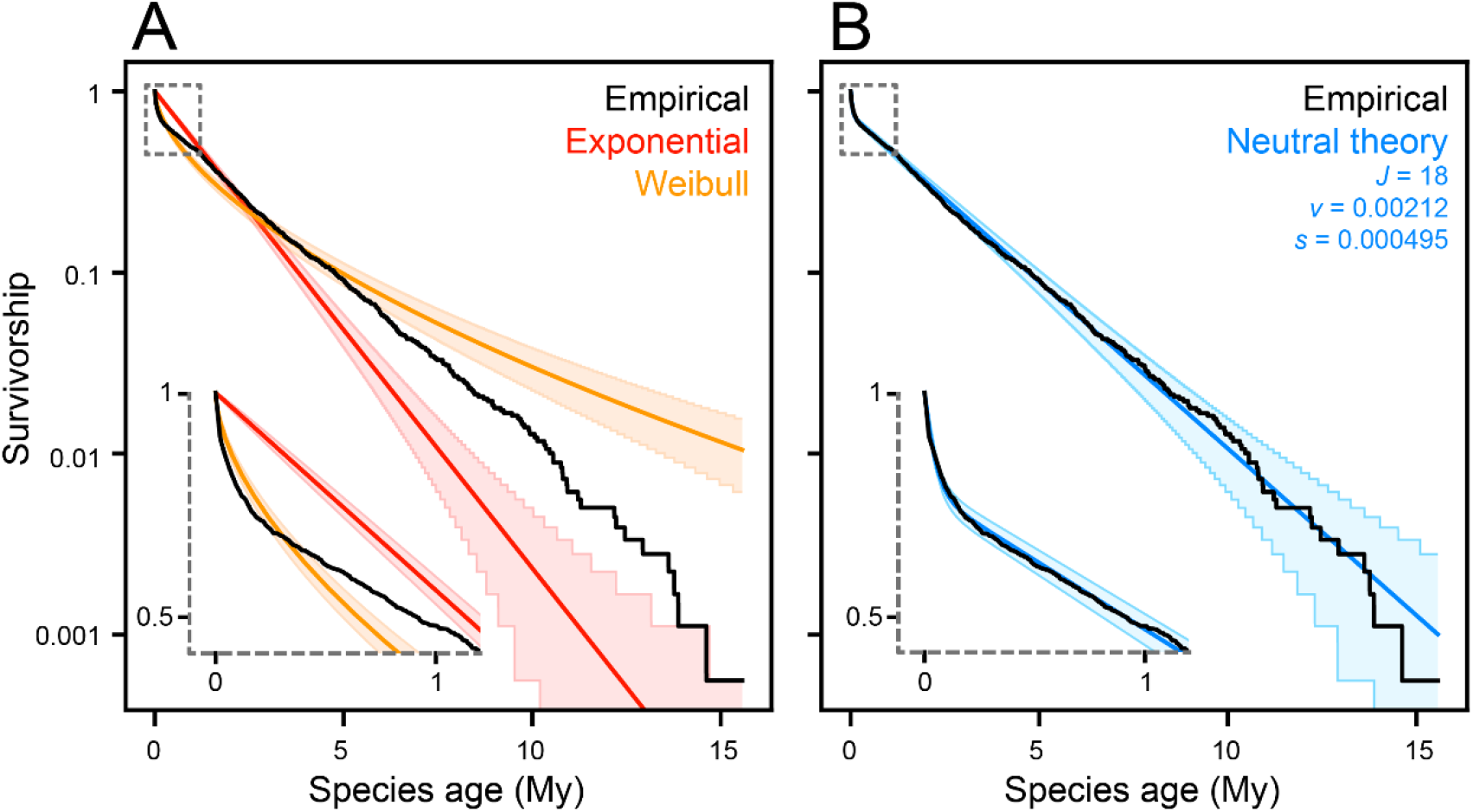
Modeling survivorship in graptoloids. Shown are empirical survivorship in Ordovician-Silurian graptoloids (black) and predictions for best fit models (colors) with 95% prediction intervals. Insets (grey dashed lines) show first million years of survivorship curve. (A) Models of age-independent (exponential) and age-dependent (Weibull) extinction. (B) Best-fit survivorship curve and associated parameter values for NT, the model overwhelmingly favored by BIC.

NT best fits demonstrate biologically interpretable differences between Ordovician and Silurian extinction regimes (Fig. 4), providing mechanistic explanations for patterns previously recognized at the descriptive level (13). The transition from Ordovician to Silurian is accompanied by a doubling of speciation rate, corresponding to higher extinction rates among old species in the Silurian (18). Likewise, inferred Ordovician community size is three times that for the Silurian, reflecting the observation that species reach constant extinction rates at a younger age in the Silurian. Likelihood ratio tests show speciation rate and community size during both the Ordovician and Silurian are significantly different from the values obtained for the entire dataset (Tables S6-S7). Finally, the cohort originating just before the Late Ordovician Mass Extinction is fit worse by NT than by a Weibull model with extinction risk increasing with species age (Fig. S3; Table S4).

**Figure 4.**
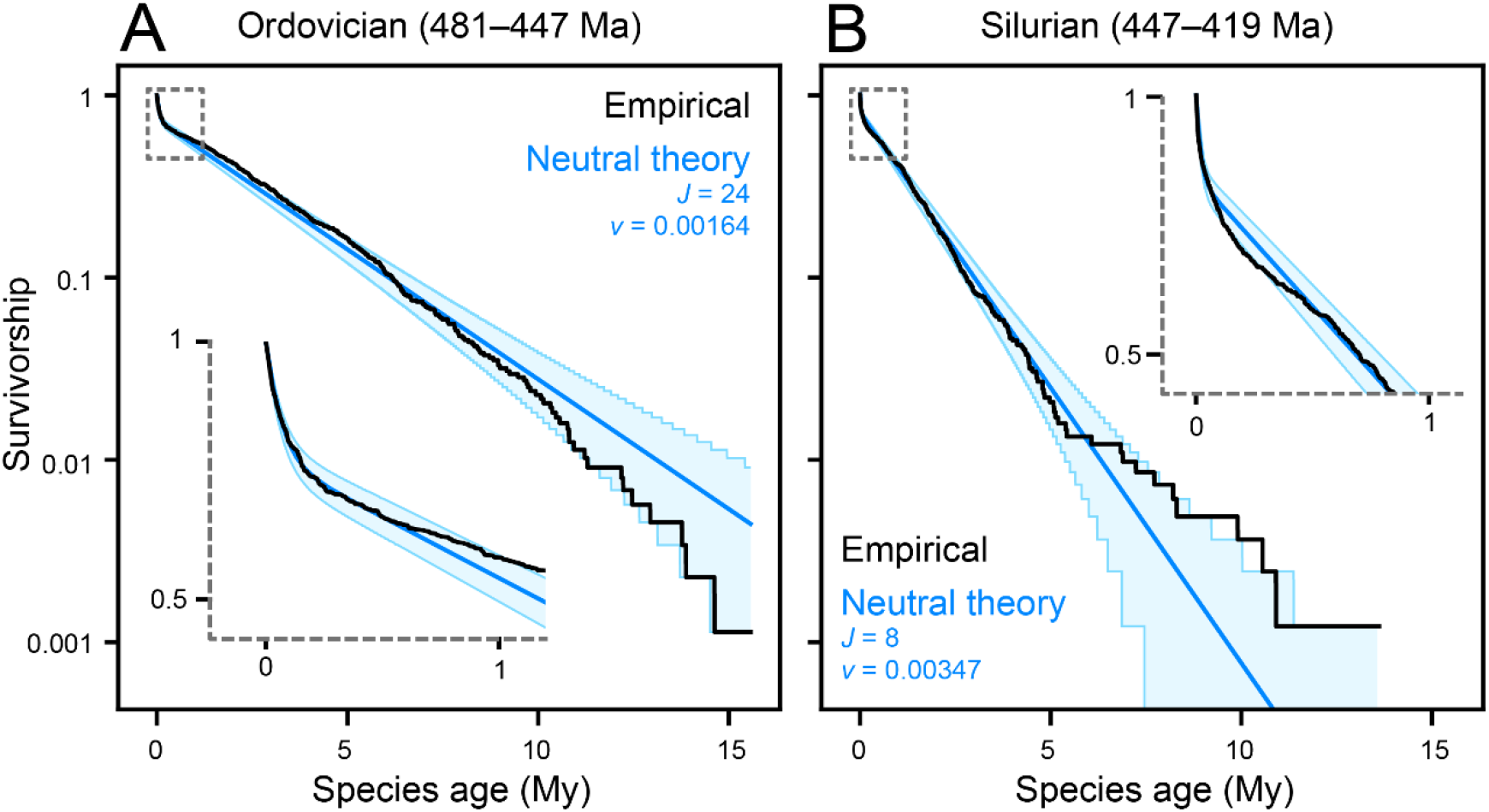
Neutral theory and changing extinction regimes. Best fit NT models and associated parameter values for (A) the Ordovician up to the late Katian (B) the late Katian–Silurian. Empirical survivorship in black, model predictions with 95% prediction intervals in blue. Insets as in Fig. 3.

## Discussion

NT succeeds as a statistical description of fossil survivorship: neutral species undergo ecological drift toward either extinction or high abundance, and the resulting survivorship curves provide exceptional fit to the graptoloid data (Fig. 3B). It remains to be seen whether this is the case for other groups of organisms. But the general predictions of NT match previous synoptic studies, which have found intense extinction among young species in most clades (6) and apparent age-independent extinction among old invertebrate taxa (5). However, the patterns of survivorship we describe might be hard to detect and study in many study systems: the ephemeral species (17) that make up the first part of the curve tend be rare and so could be missed in a less densely sampled dataset (Fig. 2C), leading to spuriously age-independent extinction (6). Ephemeral species might also have their durations artificially inflated in a less precisely calibrated dataset. Finally, reliable data at the species level are rare, and interpreting the more common genus-level analyses in terms of species-level processes is not straightforward (6, 18).

Beyond simply describing the data well—and in contrast to models like the Weibull—NT has parameters that are interpretable in terms of biological process; the theory can thus help structure hypotheses on ancient ecosystems. First, Silurian graptoloid species take less time to reach age-independent extinction, which in NT corresponds to a smaller community size (Table S3). Just as genetic drift is faster in smaller populations, ecological drift out of small starting abundance is faster in a small community (19). Small community size also explains the relatively low diversity of Silurian graptoloids (15). Second, increased extinction rates among old species in the Silurian are reflected in NT by a doubling of speciation rates. Evolutionary rate analyses have previously recovered high speciation rates among Silurian graptoloids, but it is noteworthy we reached the same conclusion without information on the timings of originations. This is because speciation and extinction rates are balanced in neutral communities at equilibrium, matching the finding from the fossil record that speciation and extinction rates are usually nearly equal (20). Geochemical and lithological data indicate the Ordovician-Silurian transition was accompanied by a suite of changes in the variability and mean state of marine abiotic environments (15, 21), and it is worth studying how these could have shaped the changes in community structure we infer. Finally, the high extinction rate among old species during the Late Ordovician Mass Extinction is completely outside the expectations of NT (Fig. 2C; Table S4). This could simply result from fitting time-homogeneous models to the results of a time-inhomogeneous process (22); for example, temporal variation in extinction intensity could distort or reverse the signal of age-dependent extinction. Alternatively, it might be due to genuine non-neutrality—that is, a role for species traits in surviving the mass extinction. Previous work suggests species originating during times of environmental upheaval were less vulnerable to extinction (23), and species arising during the mass extinction may have had this kind of advantage over older competitors. This inference also aligns with the hypothesis that mass extinctions represent departures from “background” macroevolutionary regimes, especially by threatening taxa which could otherwise resist extinction (24). Clades in NT can eventually drift to ecological fixation and become immortal; the assumptions of NT must therefore have been violated in at least some intervals in earth history, to explain how old, successful clades like graptoloids, trilobites, and ammonoids can become extinct.

Is NT a biologically reasonable model for extinction? Perhaps the most common objection to NT is not that its predictions fail but that ecological equivalence among species is demonstrably false: traits and ecological preferences vary among species in every community we can observe (25). It is tempting to speculate that planktonic filter-feeders such as graptoloids might have relatively homogeneous ecology or a limited capacity for niche differentiation, but the existence and importance of graptoloid niches is supported by apparent variation in ecological characteristics such as depth preference (26), and by repeated convergences on certain ecotypes in different graptoloid clades (27, 28). However, our success in applying NT does not imply a complete absence of ecological variation. Indeed, some ecologists have argued that locating real communities along a continuum from niche-structured to drift-structured is a more interesting goal than attempting to refute either the niche-based or neutralist worldview (29, 30). Graptoloids surely varied in their ecology, but at this scale, a good approximation of ecosystem dynamics can apparently be achieved without considering ecological differences among species (31). Moreover, the simplicity of NT is just what enables it to make comprehensive quantitative descriptions (31, 32), encompassing not just survivorship but also species abundance distributions, community turnover rates, standing richness, and occupancy trajectories. NT has been applied to community turnover (33, 34), abundance distributions (35), and diversity (36, 37) in the fossil record; further predictions remain mostly unexplored.

If apparent neutrality really is a pervasive feature of fossil communities, it has important implications for paleobiological research. For example, a goal of conservation paleobiology is explaining variation in extinction risk among fossil taxa as a function of traits (21, 38), but no universally successful predictors of extinction have been recovered apart from range size and other aspects of occupancy (39). Even body size, a “supertrait” controlling many aspects of ecology (40), has inconsistent effects on extinction risk (41, 42). This is the expectation under NT, in which the success or failure of any one species is stochastic and cannot be predicted from its traits. Instead, it is the overall pattern of extinctions and community turnover that is predictable, as in statistical physics where the chaotic movement of particles yields predictable macroscopic patterns (31). This is the opposite of the more typical picture of scaling in ecological and evolutionary systems, where predictable or deterministic population-level dynamics like Lotka-Volterra interactions (43) scale up to unpredictable macroevolutionary outcomes (3, 44).

Finally, instances of non-linear survivorship—including the graptoloid dataset used here—have been taken as refutations of RQ (5, 13, 45, 46). However, despite widespread rejection of Van Valen’s law of constant extinction [which is partly independent of RQ (47)], much of the essence of RQ survives in the neutral survivorship model. Both theories explain eco-evolutionary change as a result of zero-sum competition for control of trophic energy (48). Indeed, although RQ and NT have mostly isolated histories, they have been argued to represent distinct expressions of a more general, as yet unformulated theory (11, 49). In such a theory, innovation and extinction emerge continually as a consequence of interactions among coexisting ecological entities, even in the absence of environmental change [though abiotic variation can be incorporated into both RQ (1) and NT (50)]. The success of NT in modeling species survivorship (Figs. 3-4) thus demonstrates the continuing need for macroevolutionary theory in which biodiversity regulates itself.

## Materials and Methods

### Modeling

#### Abundance and survivorship

We derived formulae for the probability of observing any given species duration in NT given incomplete sampling. All code and data required to replicate our findings is available at https://github.com/jgsaulsbury/nt-surv. Related work has focused on ‘persistence times’ of species in neutral communities (51), oriented mainly toward power-law characterization of time to extirpation in local communities under migration, but paleontological application of this work is not straightforward. Our approach ultimately yields the probability of observing a given species duration as a function of the probabilities of a species being sampled and of going extinct in each timestep of its existence.

We first obtained a transition matrix (52) characterizing the probability of per-timestep abundance changes of a species in a neutral community defined by community size *J* and speciation rate *ν*. This approach is similar to one taken by Hubbell (7), but our approach differs from his in incorporating speciation into the transition matrix. We also do not consider migration, as that would make defining a simple abundance transition matrix impossible. Simulations indicate survivorship curves under dispersal-limitation have the same basic shape as NT without migration (Fig. S5). Nevertheless, developing NT to model survivorship in the metacommunity might yield more realistic parameter estimates, and would improve realism for global analyses such as this one (SI, Migration).

We allow one individual to be removed from the community every timestep, so a species’s abundance can change by at most one individual per timestep. Eqns. 1-3 follow Hubbell’s (7) convention for the left-hand side as P{state at time *t*+1|state at time *t*}. The probability of species *i* with abundance *N*_*i*_ increasing in abundance is

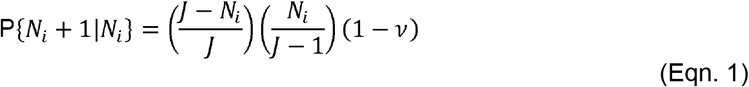

The right hand side is the probability that a death occurs in a species other than *i*, that species *i* then reproduces, and that the new individual of species *i* does not become a new species. The probability of a decrease is

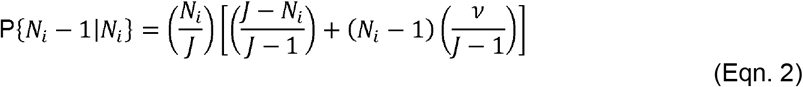

To decrease in abundance, species *i* has to lose an individual. Then, either a different species must reproduce, or else species *i* can reproduce and its offspring become a new species. The probability of undergoing no change in abundance is then

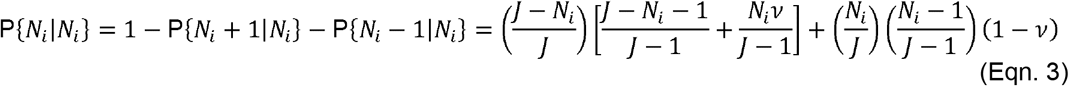

From Eqns. 1-3 we can define a *J*+1 x *J*+1 transition matrix ***M*** with entries ***M***_*j,k*_ = *P* {*N*_*i*_ *= k* | *N*_*i*_ *= j*}. Only the diagonal and first off-diagonals of ***M*** have nonzero entries, and ***M*** is absorbing at state *N*_*i*_ *=* 0. Species at maximum possible abundance *J* can still be displaced by a new species, so state *N*_*i*_ *= J* is not absorbing. A species starts with abundance 1, represented by a probability vector ***π***_0_ = {0,1,0,0,…,0} with length *J*+1. The abundance of that species at time *t* is then given by

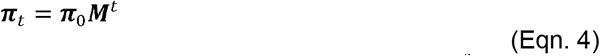

where ***M***^t^ is ***M*** to the power of *t*. The probability of being extinct by time *t* is then the 0^th^ index,*π*_*t*_[0]. Plotting 1 − *π*_*t*_ [0] against *t* yields a survivorship curve *S*(*t*), the probability that a species survives to time *t*.

To condition abundance on survival, we can define a conditional probability vector

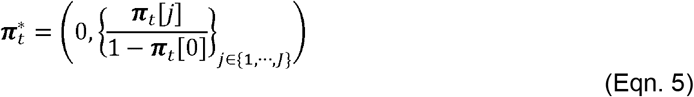

representing the probability being at some abundance at time *t*, conditional on survival to time *t*. The relationship between 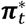 and *t* is shown in Fig. 2B. ***M*** is absorbing and has no true stationary distribution, but it has a stationary conditional distribution (53) (Fig. 2B).

Finally, we calculate the hazard function *h*(*t*), the probability that a species extant at time *t* goes extinct by time *t*+1:

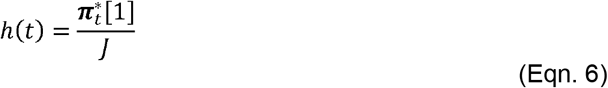

This is the probability of having a single individual, times the probability of that individual dying between time *t* and *t*+1 (cf. eqn. 2).

#### Incomplete sampling

We model incomplete sampling by giving each individual a per-timestep chance of being sampled *s*. The sampled duration of a species is the amount of time between the first and last timesteps an individual in that species is sampled. Sampling occurs at the start of a timestep. More abundant species are more likely to be detected. The probability of sampling a species at time *t* is

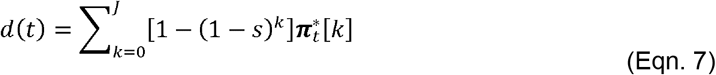

This is the probability of detecting a species with abundance *k*, times the probability of being at abundance *k*, summed over all possible *k*. The probability of sampling a species, *d*, increases with *t* until it reaches an asymptote (Fig. S1).

For the full theory (below) it will be useful to define two more terms. Let the probability of an extinction occurring between time *t* and *t*+1 be

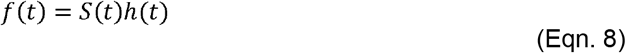

Then let the probability of *not* being detected between time 0 and *t*, inclusive, be

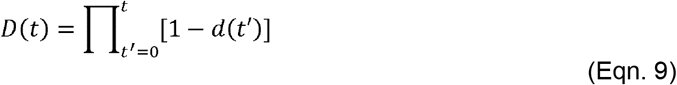

#### Full theory

We use *S, h, f, d*, and *D* to derive the probability of observing a species duration Δ*t* given incomplete sampling. This is equal to the probability of a first observation at *t* and a last observation at *t+*Δ*t*, for t between 0 and infinity. The probability for the first observation is simply

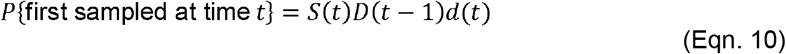

because having a first sample at time *t* requires surviving to time *t*, not being sampled before *t*, and then being sampled at time *t*. The probability for the last sampling is

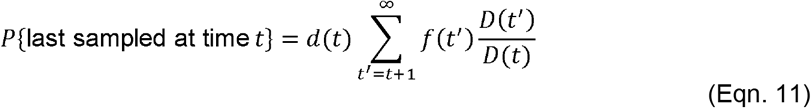

which corresponds to a final sampling at time *t*, extinction at the end of time *t*^′^, and no sampling between *t* and *t*^′^, for every possible *t′* from *t+1* to infinity. Combining these, we get

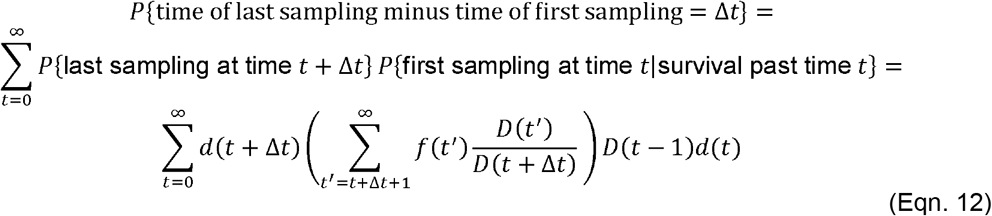

The *S(t)* term drops out in eqn. 12 because survival past the time of the first observation is implicit in eqn. 11. Eqn. 12 gives the probability of observing species durations with incomplete sampling. However, in our empirical analyses we follow (13) in excluding species with duration 0, so we condition on being at least 1 by dividing *P*{Δ*t*} by (1 – *P* {no detections} – *P* {only one detection}). Those reduce to

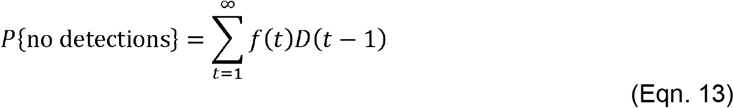

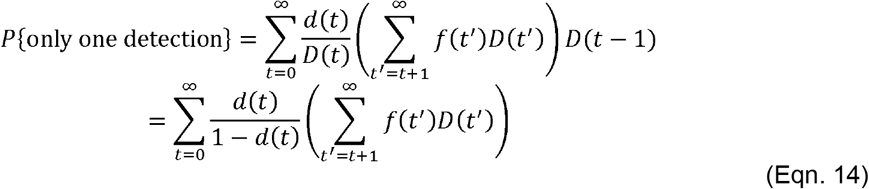

We use the resulting probability function *P*{Δ*t*} to calculate likelihoods of observing empirical durations under NT. The corresponding survivorship curve is 1 minus the cumulative distribution function for P{Δ*t*}, and this is plotted in Figs. 2-4. Equations 12-15 sum to infinity because a duration of length Δ*t* can be observed for an incompletely sampled species with any true duration >Δ*t*. However, because species rarely survive to very great durations, we can approximate P{Δ*t*} by substituting ∞ with a large number. We achieved good approximations by replacing ∞ with a value ∼1.2 times that of the largest Δ*t*.

### Dataset

We analyzed a dataset of 2041 estimated graptoloid species durations (first and last appearance times, in Ma) from (13). This dataset is a high-resolution global composite of hundreds of local stratigraphic sections recording the occurrences of graptoloid species, their position along a stratigraphic section, and the positions of independently dated events (including geomagnetic field reversals, radioisotopic dates, and stable isotopic excursions) (12, 14, 54, 55). A constrained optimization algorithm is used to find a global composite ordering of stratigraphic events (first and last appearances of species and dated events) that minimizes misfit to local sections. This ordinal composite is then “stretched” to the interval scale using local section thickness, and then to absolute time using dated events (55).

Species with estimated durations of zero (identical first and last occurrence) present a problem for survivorship analysis: it is impossible for the exponential and Weibull distributions to produce species of zero duration. These durations could conceivably be modified and included, for example by adding a very small number to their duration, but this kind of arbitrary choice could influence downstream analyses. We instead excluded the 247 species with duration 0 from our analyses. This should tend to reduce the signal of negative age-dependence in the extinction data and is therefore conservative with regard to finding support for NT (15). To explore whether our findings might be sensitive to the exceptional precision of the CONOP dataset, we conducted model fitting analyses on the main dataset rounded up to a coarser resolution of 0.05My, reaching the same conclusions (SI, Temporal resolution, Table S8).

### Analyses

NT makes predictions for species durations in units of timesteps, but the graptoloid species durations are in My. NT has been fruitfully applied with the modification that NT individuals represent subpopulations (56); we used this approach to model survivorship in NT because modeling realistic community sizes and timescales was computationally infeasible. This approach may also be preferable because, similar to Allen & Savage’s (57) extension of NT, new species begin as a single subpopulation rather than as a single individual. We define each NT individual as a subpopulation with an expected lifespan of 5000 years. The decision to use this value to convert between geologic time and model time is pragmatic rather than theoretically grounded. However, the same survivorship curves in geologic time can be obtained under a variety of values for this conversion factor (Fig. S2). Thus, the survivorship curve appears robust to changes in the defined scale of a subpopulation. Furthermore, parameters have the same effect on the resulting survivorship curve regardless of conversion factor. The slope of the second, age-independent phase of the survivorship curve is always directly proportional to *v* because the per-timestep extinction probability of old species is *v* divided by *J*. Likewise, the characteristic timescale for the approach to stationarity in NT [tau in (31)] is proportional to community size, so, all else equal, an increase in J will yield a proportional increase in the time to reach constant extinction.

We convert species durations from millions of years to expected lifespans as 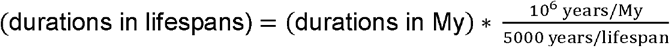. In NT, the expected lifespan of an individual is *J* timesteps. So, we convert empirical durations from lifespans to NT timesteps using (durations in timesteps) = (durations in lifespans)\**J*, rounding up to the nearest integer.

We found maximum likelihood estimates (SI, Fitting NT) for *J, v*, and *s* for the entire dataset of 1794 species and for subsets (see Introduction). For comparison, we used the *R* package *fitdistrplus* (58) to fit exponential and Weibull models to the data, and compared models using Akaike weights. To assess model adequacy, we calculated Kolmogorov-Smirnov statistics (the maximum absolute distance between modeled and empirical survivorship) for NT model fits and compared these with critical values for the one-sample Kolmogorov-Smirnov test. We compared models with BIC, which is designed to favor the model with the highest average likelihood across parameter space and penalizes overfitting more aggressively than the more commonly used AIC (59). AIC values are provided for comparison.

Finally, we generated prediction intervals for all survivorship curves using the binomial distribution. For a sample of size *n*, the number of species surviving at least to age *t* is given by the binomial distribution *B*(*n,S*(*t*)) for survivorship function *S(t)*. The 2.5^th^ and 97.5^th^ percentiles of this distribution yield a 95% prediction interval.

## Acknowledgments

We thank Tomasz Baumiller, Seth Finnegan, Kelly Matsunaga, and Carolann Schack for suggestions and discussions that improved this work, as well as Kjetil Voje, Steve Holland, John Alroy, and an anonymous reviewer for thoughtful comments and criticism on earlier drafts. We especially thank James Crampton, Peter Sadler, and the late Roger Cooper for the dataset of graptoloid species durations.

## Notes

https://github.com/jgsaulsbury/nt-surv

### Competing Interest Statement

The authors have declared no competing interest.

