## Supplementary material for "Age-dependent extinction and the neutral theory of biodiversity": SI

Supporting Information Text

**Fitting NT.** The likelihood of a neutral theory (NT) model for a set of species durations is the product of all P{Δt} for every Δt in the dataset. Likelihood surfaces for NT survivorship models appear to be convex with respect to community size, speciation rate, and sampling rate, so we found parameter values that maximized likelihood through a simple hill-climbing approach. This is straightforward for speciation rate and sampling rate, but problematic for community size because the empirical durations are converted to units of timesteps using a given value of community size. For example, a species duration of length 100 lifespans would represent 500 timesteps given *J*=5 and 1000 timesteps given *J*=10. Likelihoods are only comparable with respect to the same data. So, for fitting values of *J*, we used species durations in lifespans (rounded up to the nearest integer) which are the same regardless of *J*. This required applying a cumulation process to the likelihood function P{Δt} because durations in different numbers of timesteps were rounded up to the nearest lifespan. For example, if J=5, P{2 lifespans} = P{6 timesteps}+ P{7 timesteps}+ P{8 timesteps}+ P{9 timesteps}+ P{10 timesteps}. Once we found parameter values that maximized fit for the entire dataset as well as for Ordovician, Silurian, and Late Ordovician Mass Extinction subsets, all other analyses (including AIC comparisons and all figures) were done using durations in timesteps, not lifespans.

**Temporal resolution**. Ages of first and last for species in the CONOP database are given with high precision (4 decimal places, for a resolution of 100 years). Time between event levels in the CONOP dataset varies from 10^3^ to 10^6^ years, with most between 10^4^ and 10^5^ and an average of 0.037My. To assess how sensitive our findings are to the precision of the data, we rounded all durations up to the nearest 0.05 My and reran model fitting analyses for the full dataset. The conclusions are not affected (Table S8). NT still receives decisive support from AIC, with a collective AIC of less than 10^-21^ for exponential and Weibull models. BIC also supports NT, with ΔBIC = 91 for the next-best model (Weibull). We conclude that our model-fitting results are not being driven by the exceptionally short (<0.01Myr) durations of some of the graptolite species in the dataset, or more generally by the high precision with which the data are reported. We note, however, that these high-precision durations, including the exceptionally short ones, are not in some way an artifact of the CONOP procedure, but are instead based on real stratigraphic measurements from local sections. We therefore run all other analyses on the dataset in the original resolution.

**Migration.** The analysis presented in the main text uses a version of NT without a migration term. This simplification allows us to formulate abundance dynamics in a transition matrix, which in turn allows us to calculate likelihoods, but is arguably unrealistic for the global dataset we work on. We explore the effects of this simplifying assumption by using simulations to assess survivorship under dispersal limitation. We set up the metacommunity as a square grid of 64 local communities, each of size *J* = 8, for a metacommunity size of *J_M_* = 512. The rules of demographic change are the same as in the local community (Fig. 1, main text), except that, whenever the replacement for a lost individual would be drawn from its local community, there is an *m* chance it will instead be drawn from the metacommunity. Migrants to a local community are drawn from the Moore neighborhood of that community (the eight communities surrounding it in a grid). We simulated survivorship with complete sampling for 50,000 timesteps in a metacommunity with *v* = 0.005. We ran simulations with migration rates *m* of 0, 0.02, 0.1, and 0.5. Results are plotted alongside analytically derived survivorship curves for a local community of size *J* and one of size *J_M_* (Fig. S5). Results are similar to the survivorship curves from the main analysis: concave-up and converging on constant extinction rate among old species. Survivorship in the metacommunity is the same as in the local community when migration rate *m* = 0, and when *m* approaches 1, survivorship approaches that in a local community of size *J_M_* (the number of individuals in the metacommunity). Intermediate migration rates interpolate between these two extremes. Because the local community model is a special case of the metacommunity model, the best metacommunity model must be at least as good as the best local community model.

Considering survivorship in a metacommunity may also help explain parameter estimates. In particular, fitted values of community size *J* seem low, even considering the subpopulation substitution: assuming a log-linear inverse relationship between conversion factor (see Methods) and fitted *J* (Fig. S2) and extrapolating to realistic lifespans for individuals yields community sizes in the hundreds of thousands. The global graptoloid community was probably much larger than this on average, but J in the millions or billions would predict a longer wait time to reach the age-independent phase of the survivorship curve than the observed one of about ~200ky (Fig. 3). We expect considering survivorship in the metacommunity will help resolve this misfit: even in a very large metacommunity, a low migration rate can result in species survivorship that closely resembles that expected for a small community (Fig. S5). In other words, when the global biota is subdivided into local communities linked by rare migration, it is the size of the local community that mainly determines expected survivorship, regardless of global community size. Another possibility, not mutually exclusive with the first one, is that intermittent bottlenecks lead to an “effective community size (1)” much lower than the average size. Survivorship in the metacommunity will be more challenging to model but would help achieve plausible parameter estimates, while simultaneously generating new kinds of biogeographic predictions.

**Data and code.** All code and data required to replicate our results is available at https://github.com/jgsaulsbury/nt-surv.

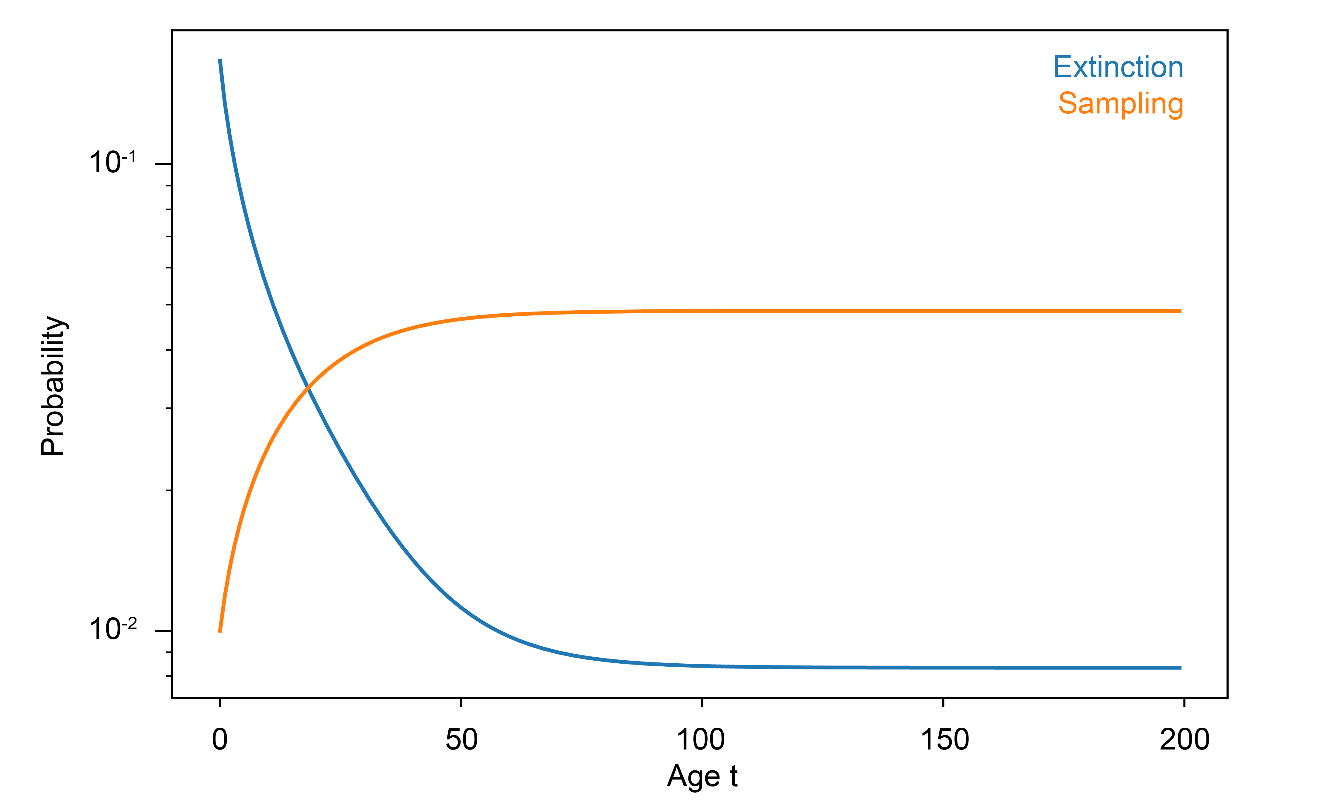

Fig. S1. Per-timestep sampling and extinction probabilities versus the age of a species surviving to time t. Community parameters same as those in main text Fig. 2A-B. Per-species per-time sampling probability increases in a similar way as expected abundance, while per-timestep extinction probability decays from 1/*J* to ν/*J*, where *J* is community size and *ν* is speciation rate.

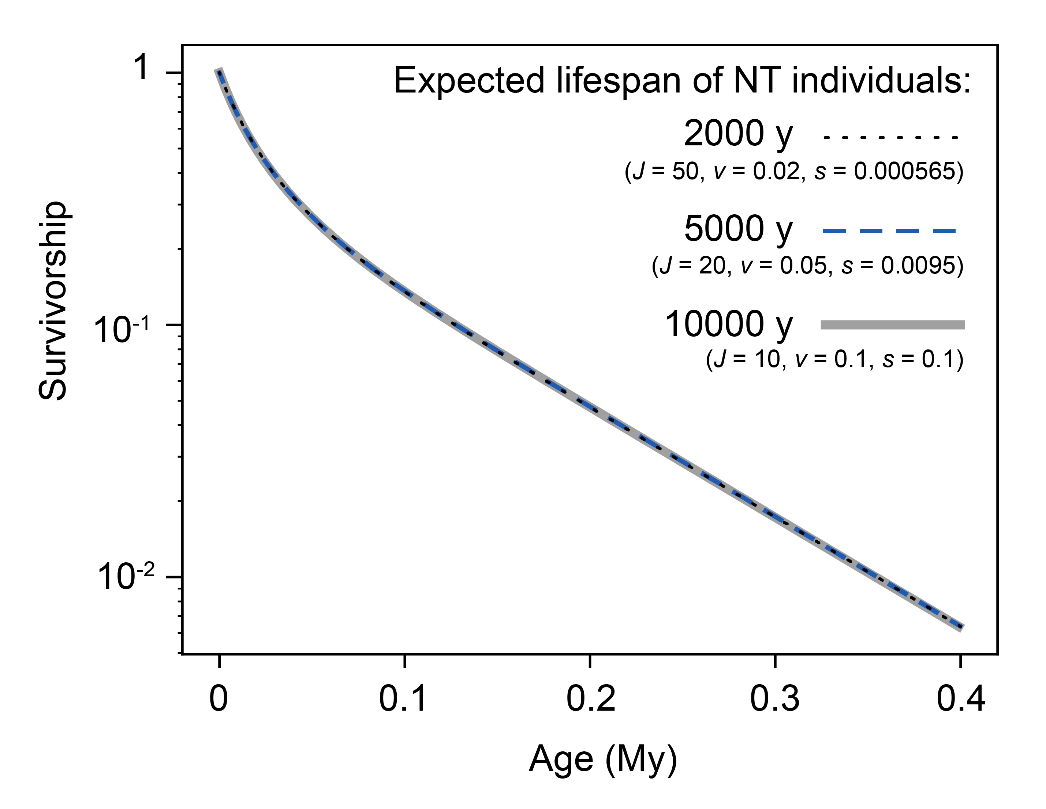

Fig. S2. Identical survivorship curves obtained under three different choices of the conversion factor between model time and geologic time. The choice of conversion factor is arbitrary, but our findings are robust to different choices for this conversion. *J* is community size, *v* is speciation rate, and *s* is per-individual per-timestep sampling rate.

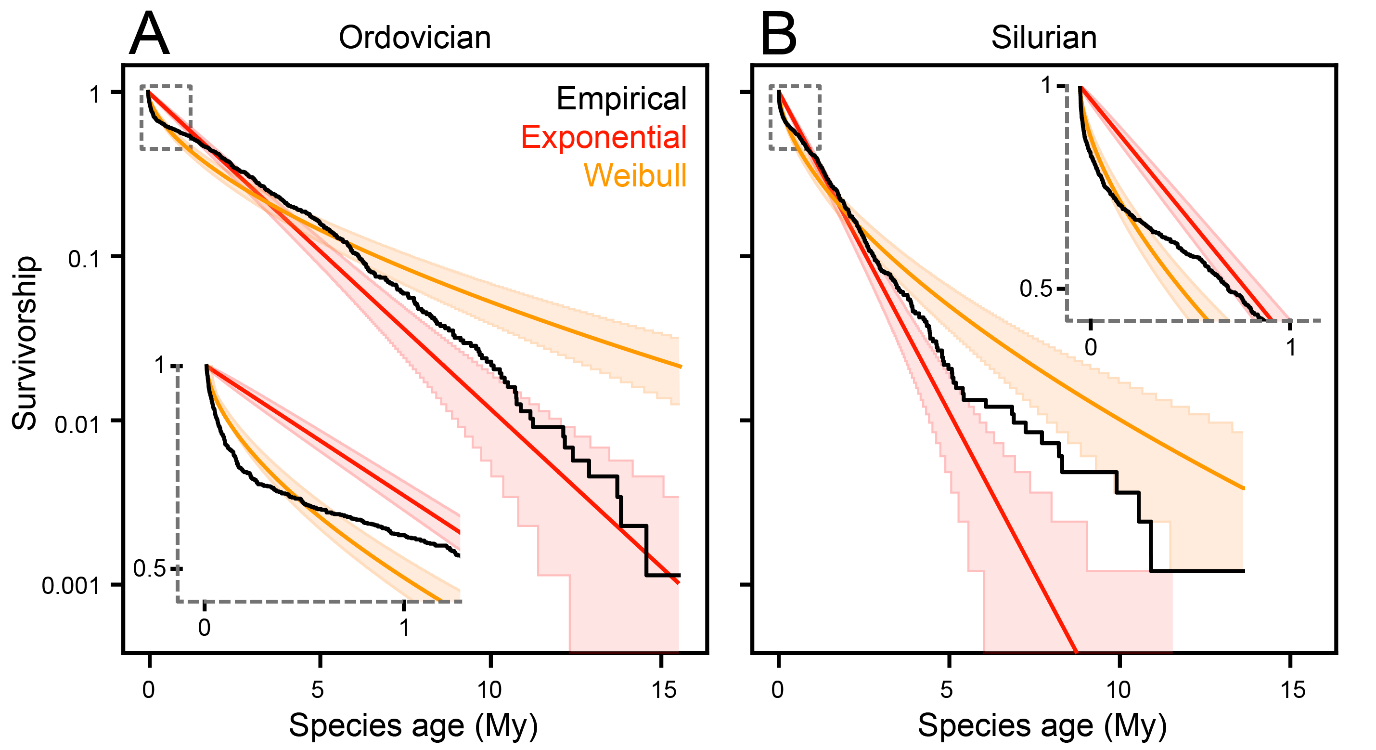

Fig. S3. Empirical survivorship curves (black) with best-fit exponential and Weibull survivorship models for subsets of the data. (A) Ordovician. (B) Silurian.

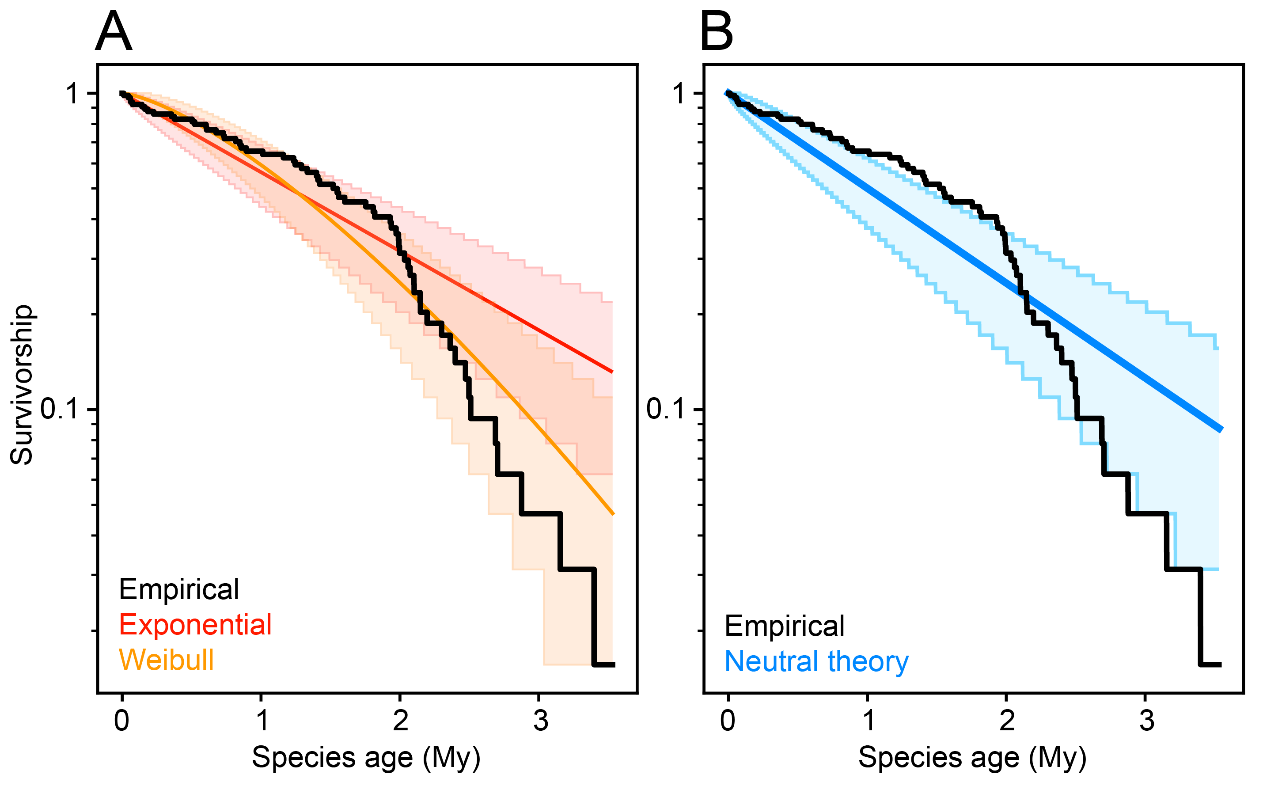

**Fig. S4.** Empirical survivorship curve for the 64 species originating immediately before the Late Ordovician Mass Extinction (450–448 Ma, black), along with best-fit survivorship models. This cohort is best fit by a Weibull model with extinction probability increasing with species age (Table S4). (A) Weibull (orange) and exponential (red) models fit to the data. (B) The same data, with NT best-fit model.

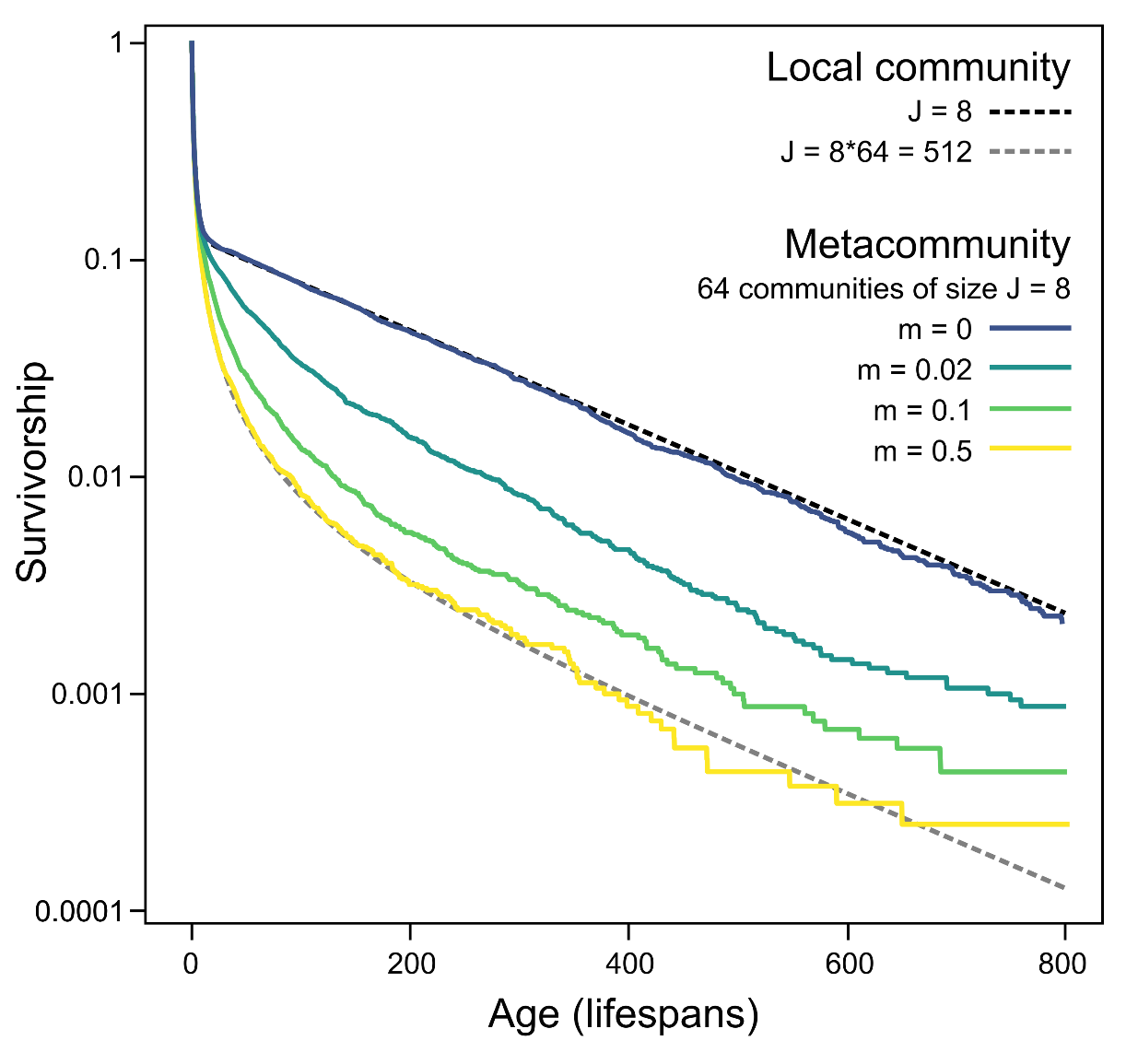

**Fig. S5.** Survivorship in a metacommunity of 64 linked local communities of size *J* = 8. Changing migration rate *m* interpolates between the survivorship curve expected in a single local community of size *J*, and one in a local community of size *J_M_* = 512.

**Table S1.** Best-fitting models, full dataset. Also shown are log-likelihoods and AIC and BIC scores. N = 1794.

**Table S2.** Best-fitting models, Ordovician (minus the Hirnantian and late Katian) subset (481-447 Ma). N = 880.

| **Model** | **Scale** | **Shape** | **J** | **nu** | **s** | **loglik** | **free parameters** | **AIC** | **dAIC** | **Akaike weight** | **BIC** | **dBIC** |
| --- | --- | --- | --- | --- | --- | --- | --- | --- | --- | --- | --- | --- |
| Exponential | 5939.265 | - | - | - | - | -17465.8 | 1 | 34933.5 | 1078.836 | 5.42E-235 | 34938.99 | 1067.852 |
| Weibull | 4363.482 | 0.594194 | - | - | - | -17010.5 | 2 | 34025.02 | 170.356 | 1.02E-37 | 34036 | 164.8638 |
| NT | - | - | 18 | 0.00212 | 0.000495 | -16924.3 | 3 | 33854.66 | 0 | ~1 | 33871.14 | 0 |

| **Model** | **Scale** | **Shape** | **J** | **nu** | **s** | **loglik** | **free parameters** | **AIC** | **dAIC** | **Akaike weight** | **BIC** | **dBIC** |
| --- | --- | --- | --- | --- | --- | --- | --- | --- | --- | --- | --- | --- |
| Exponential | 10840.16 | - | - | - | - | -9084.2 | 1 | 18170.4 | 548.112 | 9.53E-120 | 18175.9 | 537.1276 |
| Weibull | 8170.766 | 0.608275 | - | - | - | -8891.43 | 2 | 17786.86 | 164.564 | 1.84E-36 | 17797.84 | 159.0718 |
| NT | - | - | 24 | 0.00164 | 0.00021 | -8808.15 | 3 | 17622.29 | 0 | ~1 | 17638.77 | 0 |

**Table S3.** Best-fitting models, Silurian (plus the Hirnantian and late Katian) subset (447-419 Ma). N = 826.

| **Model** | **Scale** | **Shape** | **J** | **nu** | **s** | **loglik** | **free parameters** | **AIC** | **dAIC** | **Akaike weight** | **BIC** | **dBIC** |
| --- | --- | --- | --- | --- | --- | --- | --- | --- | --- | --- | --- | --- |
| Exponential | 1830.487 | - | - | - | - | -7031.27 | 1 | 14064.53 | 449.1374 | 2.96E-98 | 14070.02 | 438.153 |
| Weibull | 1346.356 | 0.618401 | - | - | - | -6858.35 | 2 | 13720.69 | 105.2994 | 1.36E-23 | 13731.68 | 99.80723 |
| NT | - | - | 8 | 0.00348 | 0.0047 | -6804.7 | 3 | 13615.39 | 0 | ~1 | 13631.87 | 0 |

**Table S4.** Best-fitting models, cohort originating immediately before Late Ordovician Mass Extinction (450-448 Ma). N = 64.

| **Model** | **Scale** | **Shape** | **J** | **nu** | **s** | **loglik** | **free parameters** | **AIC** | **dAIC** | **Akaike weight** | **BIC** | **dBIC** |
| --- | --- | --- | --- | --- | --- | --- | --- | --- | --- | --- | --- | --- |
| Exponential | 3479.546 | - | - | - | - | -576.036 | 1 | 1154.073 | 8.2862 | 0.015549 | 1159.565 | 2.793997 |
| Weibull | 3181.64 | 1.401756 | - | - | - | -570.893 | 2 | 1145.787 | 0 | 0.979548 | 1156.771 | 0 |
| NT | - | - | 10 | 0.00345 | 0.0001 | -575.191 | 3 | 1156.381 | 10.59462 | 0.004903 | 1172.858 | 16.08682 |

**Table S5.** Kolmogorov-Smirnov tests. Parameters same as in best fit models (Tables S1-S3).

|  |  | **KS critical value** | | | | **Exponential** | | **Weibull** | | **NT** | |
| --- | --- | --- | --- | --- | --- | --- | --- | --- | --- | --- | --- |
| **Dataset** | **N** | **α = 0.05** | **α = 0.02** | **α = 0.01** | **α = 0.001** | **KS statistic** | **p-value** | **KS statistic** | **p-value** | **KS statistic** | **p-value** |
| Entire dataset | 1794 | 0.032064209 | 0.035825928 | 0.038427471 | 0.046026223 | 0.212011 | p < 0.001 | 0.0867425 | p < 0.001 | 0.034443512 | 0.05 > p > 0.02 |
| Ordovician subset | 880 | 0.045781542 | 0.051152555 | 0.054867059 | 0.06571662 | 0.2234506 | p < 0.001 | 0.0988966 | p < 0.001 | 0.04648132 | 0.05 > p > 0.02 |
| Silurian subset | 826 | 0.047254343 | 0.052798142 | 0.056632143 | 0.067830737 | 0.1967645 | p < 0.001 | 0.0935247 | p < 0.001 | 0.042874842 | p > 0.05 |

**Table S6.** Likelihood ratio tests, Ordovician subset. The best-fitting model (top row) is compared with one in which the community size *J* (middle row) and the speciation rate *v* (bottom row) are set to the values obtained from fitting NT to the full dataset.

| **model** | ***J*** | ***nu*** | ***s*** | **loglik** | **LRT statistic** |
| --- | --- | --- | --- | --- | --- |
| unconstrained | 24 | 3.28E-07 | 0.00021 | -6011.41 | - |
| *J* set to 18 | 18 | 3.22E-07 | 0.00045 | -6016.13 | 9.436123 |
| *nu* set to 0.000495 | 22 | 4.24E-07 | 0.00025 | -6039.05 | 55.28126 |

**Table S7.** Likelihood ratio tests, Silurian subset. The best-fitting model (top row) is compared with one in which the community size *J* (middle row) and the speciation rate *v* (bottom row) are set to the values obtained from fitting NT to the full dataset.

| **model** | ***J*** | ***nu*** | ***s*** | **loglik** | **LRT statistic** |
| --- | --- | --- | --- | --- | --- |
| unconstrained | 8 | 6.95E-07 | 0.0047 | -5090.614 | - |
| *J* set to 18 | 18 | 6.44E-07 | 0.00052 | -5098.824 | 16.420 |
| *v* set to 0.000495 | 15 | 4.24E-07 | 0.0009 | -5142.745 | 104.263 |

**Table S8.** Model fitting results obtained when binning the data to 50ky resolution (c.f. Table S1)

| **Model** | **Scale** | **Shape** | **J** | **nu** | **S** | **loglik** | **free parameters** | **AIC** | **dAIC** | **relative likelihood** | **Akaike weight** | **BIC** | **dBIC** |
| --- | --- | --- | --- | --- | --- | --- | --- | --- | --- | --- | --- | --- | --- |
| Exponential | 6149.022 | - | - | - | - | -17492.4 | 1 | 34986.78 | 498.4407 | 5.82E-109 | 5.82E-109 | 34992.27 | 487.4563 |
| Weibull | 5058.831 | 0.707407 | - | - | - | -17290.6 | 2 | 34585.16 | 96.82066 | 9.45E-22 | 9.45E-22 | 34596.14 | 91.32846 |
| NT | - | - | 18 | 4.56E-07 | 0.00037 | -17241.2 | 3 | 34488.34 | 0 | 1 | ~1 | 34504.82 | 0 |
